# A fast fish swimming protocol that provides similar insights as critical sustained swimming speed

**DOI:** 10.1101/2024.04.10.588974

**Authors:** Stephanie M. Bamford, Frank Seebacher

## Abstract

Performance measures are an important tool to assess the impact of environmental change on animals. In fish, performance is often measured as critical sustained swimming speed (U_crit_), which reflects individual physiological capacities. A drawback of U_crit_ is that trials are relatively long (∼30-75 min). U_crit_ is therefore not suitable for repeated measurements because of the potential for training effects, long recovery periods, and low throughput. Here we test a shorter (∼4-5 min) protocol, “U_crit_ fast” (U_Cfast_) in zebrafish (*Danio rerio*). We show that U_Cfast_ and U_crit_ have similar, significant repeatabilities within individuals. Unlike U_crit_, repeated U_Cfast_ trials do not elicit a training effect. Both U_Cfast_ and U_crit_ provide the same insights into thermal acclimation, and both provide similar estimates of individual acclimation capacity in doubly acclimated fish. We propose that U_Cfast_ is a valid substitute for U_crit_ particularly when higher throughput and repeated measures are necessary.

## Introduction

Performance may be defined as the value of a trait that is closely related to fitness, such as metabolic and locomotor traits (Husak et al., 2006; Le Galliard et al., 2004). The capacity to move fast enough to catch prey or long enough to disperse to new habitats will determine the success of animals in their natural environment (Denton et al., 2017; Domenici et al., 2019; Wu and Seebacher, 2022). Locomotor performance is therefore closely related to fitness, but it may also be costly so that there are trade-offs that determine individual fitness (Husak and Lailvaux, 2022). Environmental conditions can impact these responses and measures of locomotor performance are important to assess the impacts of environmental change on animals (Domenici et al., 2019; Killen et al., 2017; Killen et al., 2021).

Locomotor performance is a widely used whole-animal performance measures in the literature, particularly in fish (Wu and Seebacher, 2022). The most common modes of locomotor performance measured are sprint speed and critical sustained swimming speed (U_crit_)(Domenici and Blake, 1997; Kolok, 1999). Sprint speed, often measured as an escape response, is linked to predator-prey interactions where it can determine escape success (Domenici et al., 2019). Sprints are classified physiologically as high-intensity movements that are powered anaerobically and last for less than 30 seconds (Burgomaster et al., 2008; Marras et al., 2013), although escape responses in fish are much shorter than this. In contrast, U_crit_ protocols are much longer (>30 min depending on context), and measure combined anaerobic and aerobic locomotor capacities (Marras et al., 2013; Plaut, 2001; Svendsen et al., 2010). U_crit_ tests in fish are similar to treadmill tests in humans and typically use swimming flumes in which fish swim at gradually stepped-up water flow speeds until a speed is reached when animals can no hold their position in the water flow (Plaut, 2001). U_crit_ is then calculated from the duration of each speed step and the time fish were able to swim at their final step. Actual flow speeds and the durations of steps vary between protocols and for different species. In a typical protocol for zebrafish, water flow steps are 5-10 min in duration, and fish start the trial swimming at approximately 1-3 body lengths s^-1^ (0.03-0.1 ms^-1^), and at each step flow rate is increased by 1-2 body lengths s^-1^ (Seebacher and Bamford, 2024; Thambithurai et al., 2019). The total protocol lasts somewhere between 30 and 75 min depending on size, test temperature, and thermal history of fish. U_crit_ depends on the combined capacities of the cardiovascular system, energy metabolism, and neuromuscular function and is therefore an excellent indicator of overall physiological performance.

The length of U_crit_ trials are a drawback both in terms of the exercise intervention for the fish, and the time taken to complete a single run. Hence, the method has relatively low throughput, and does not lend itself to repeated measures because fish need a relatively long time to recover from the exhaustive exercise (Zhang et al., 2018) and there is a likelihood of a training effect (Hammer, 1995; Simmonds and Seebacher, 2017). We therefore aimed to test a modified U_crit_ protocol (U_Cfast_) that reduces swimming time to around 5 min. The protocol reduces the duration of each speed step while keeping other parameters (e.g. magnitude of steps) the same. The rationale for the U_Cfast_ protocol is that anaerobically fuelled locomotion that relies primarily on fast muscle fibres is of only very short duration (<<30s). Hence, a locomotor trial even lasting 5 min would rely on a combination of fast and slow muscle fibres and aerobic and anaerobic ATP production. We hypothesised therefore that a ∼5 min protocol should give similar responses as the much longer U_crit_ protocol. We tested this hypothesis in zebrafish (*Danio rerio*), testing repeatability of U_crit_ and U_Cfast_, and implementing acclimation treatments that provided a range of different contexts for comparison between the two approaches.

## Materials and methods

### Study animals

Adult zebrafish of mixed sex (*Danio rerio*; mean standard length = 3.28 ± 0.02 [s.e.] cm; mean mass = 0.56 ± 0.014 [s.e.] g) were obtained from a commercial supplier (Livefish, Bundaberg, Australia). After arrival fish were dispersed across five tanks (0.65 x 0.28 x 0.32 m) filled with filtered, aged water at 24°C until use in experiments after 1-2 weeks. During this holding period and in all experiments, water temperatures were maintained in a temperature-controlled room and each tank contained a biological filter and aerator. Fish were fed daily with pellet food (O.range WEAN 2/4, Primo Aquaculture, Narangba QLD, Australia) supplemented with live *Artemia* twice per week. All procedures had the approval of the University of Sydney Animal Ethics Committee (approval #2021/1932).

### Swimming performance

We measured critical sustained swimming speed (U_crit_) according to published protocols (Seebacher et al., 2015) in cylindrical, clear plastic (Perspex) flumes (150 mm length, 26 mm internal diameter) tightly fitted over the intake end of a submersible inline pump (12V DC, iL500, Rule, Hertfordshire, UK). A plastic grid separated the flume from the pump, and a bundle of hollow straws at the inlet helped to maintain laminar flow. Flumes were submerged in plastic tanks (0.65 x 0.28 x 0.32 m), and we used a variable DC power source (NP9615; Manson Engineering Industrial, Hong Kong, China) to adjust the flow speed. Flow speed was measured in real-time during swimming trials using a flow meter (DigiFlow 6710 M, Savant Electronics, Taichung, Taiwan) attached to the outlet of the pump. Fish swam at an initial flow rate of 0.1 m/s for 10 min followed by an increase in flow speed of 0.06 m/s every 10 min until the fish could no longer hold their position in the water column. The first time fish fell back to the plastic grid, flow was stopped for 10 s after which the previous flow rate was resumed. The trial was stopped when fish fell back to the grid a second time.

The fast U_crit_ protocol (U_Cfast_) we tested here was identical to the U_crit_ protocol described above, but the durations of each speed increment was shortened and the second increment was used as the starting speed. Hence, fish swam at an initial speed of 0.16 m s^-1^ for 1 min, and speed was then increased by 0.06 ms^-1^ every 1 min. At 24°C (during repeatability tests, see below), the mean duration of U_crit_ trials was 49.6 ± 0.6 [s.e.] min, and of U_Cfast_ trials it was 4.7 ± 0.08 [s.e.] min.

### Repeatability

We repeatedly measured U_crit_ and U_Cfast_ in the same individuals (n = 24 fish), 1, 2, 3, and 8 days apart to determine whether swimming performance measures reflect consistent phenotypes. Trials were conducted at 24°C, and fish were dispersed across six glass tanks (0.30 x 0.18 x 0.20 m) between trials. Between swimming trials fish were kept in perforated cylindrical baskets (1 l volume) within their home tanks so that we could follow individuals while still permitting visual and olfactory contact between fish (Seebacher et al., 2015). U_crit_ and U_Cfast_ were measured 24 h apart in each fish, and the experiment was conducted in two blocks with n = 12 fish in each.

### Acclimation capacity

We measured acclimation and individual acclimation capacity (Seebacher *et al*. 2015) of fish (n = 28) by acclimating each individual to warm and cold conditions sequentially. As above, individual fish were kept in perforated cylindrical plastic containers (1 l volume) during the acclimation period. Each fish was acclimated twice, with half the fish randomly assigned to the cool (20°C) acclimation treatment first for 3-4 weeks, and then to the warm (28°C) acclimation treatment for 3-4 weeks, and the other half to the opposite order to dilute any order effects. To reach the respective acclimation temperatures, water temperatures were changed gradually by 3-4°C per day over two days. Acclimation temperatures were well within the natural range of temperatures experienced by zebrafish (López-Olmeda and Sánchez-Vázquez, 2011). During acclimation, fish were dispersed across four glass tanks (0.30 x 0.18 x 0.20 m) within each treatment. After each acclimation period, we measured U_crit_ and U_Cfast_ at 20°C and 28°C acute test temperatures in each individual. Fish were given at least 24 h to recover after each swimming performance measure. After each acclimation treatment, we photographed each fish to determine body length (using ImageJ software, National Institute of Health, USA), and we weighed each fish on an electronic balance.

Acclimating each individual to both temperatures allowed us to quantify phenotypic plasticity of each individual, expressed as an acclimation capacity index (Seebacher et al., 2015):

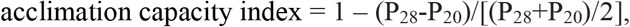

where P_28_ is the U_crit_ of a fish that is acclimated to 28°C and measured at 28°C acute test temperature, while P_20_ is the equivalent measure at 20°C. The acclimation capacity index indicates relative thermal compensation (the ability to maintain constant performance across thermal conditions) by contrasting the difference between P_20_ and P_28_. Acclimation capacity approaches 1 as P_20_ approaches P_28_, and decreases as the difference between P_28_ and P_20_ increases. The index could also be expressed in the opposite direction as P_20_ – P_28_, but fish tended to perform better in warm conditions due to the thermodynamic depression caused by cold temperatures. If a fish over-compensated for low temperatures and P_20_ > P_28_, the index will exceed 1. The index is based on the difference between P_20_ and P_28_ normalised to their mean, and is therefore a dimensionless number that is independent from the absolute values of P_20_ and P_28_.

### Statistical analysis

We analysed repeatability of swimming performance of individuals between days of measurements in the R package rptR (Stoffel et al., 2017). Repeatability (R) represents the fraction of the total phenotypic variance in the population that can be attributed to individual identities, and significance is estimated by permutational p-values (Stoffel et al., 2017). We used permutational analyses of variance in the R package lmPerm (Wheeler and Torchiano, 2016) for the remainder of the analyses. Permutational analyses are advantageous because the data per se are used for analysis and no assumptions about underlying distributions are necessary; statistical results are given as permutational p-values (Drummond and Vowler, 2012; Ludbrook and Dudley, 1998). Differences in mean performance between repeated days of swimming were analysed with day as the fixed factor and individual ID as a random factor. In analyses of acclimation responses, acclimation temperatures, acute test temperatures, and swimming performance type (U_crit_ or U_Cfast_) were fixed factors. Swimming performance was analysed in m s^-1^ with standard length of fish as a co-variate, and we used individual ID as a random variable to account for repeated measures of individual fish. Individual acclimation capacity was analysed with swimming performance type as fixed factor and individual ID as random factor.

## Results and Discussion

### Repeatability

Both U_crit_ (R = 0.79 [95% confidence interval = 0.63-0.87]; p < 0.001) and U_fast_ (R = 0.69 [95% CI = 0.51-0.81]; p < 0.001) were significantly repeatable (Fig. 1A, B). The broad overlap of confidence intervals indicates that repeatability did not differ between the two measures of swimming performance. Note that mean U_crit_ increased with repeated swimming (p = 0.0018; Fig. 1C) but U_Cfast_ did not (p = 0.98, Fig. 1C). These data indicate that either measure is a consistent representation of the intrinsic physiological performance of individual fish. However, the increase in performance following repeated U_crit_ trials is likely to represent a training effect that does not occur with the less intense exercise intervention of U_Cfast_ trials.

**Figure 1.**
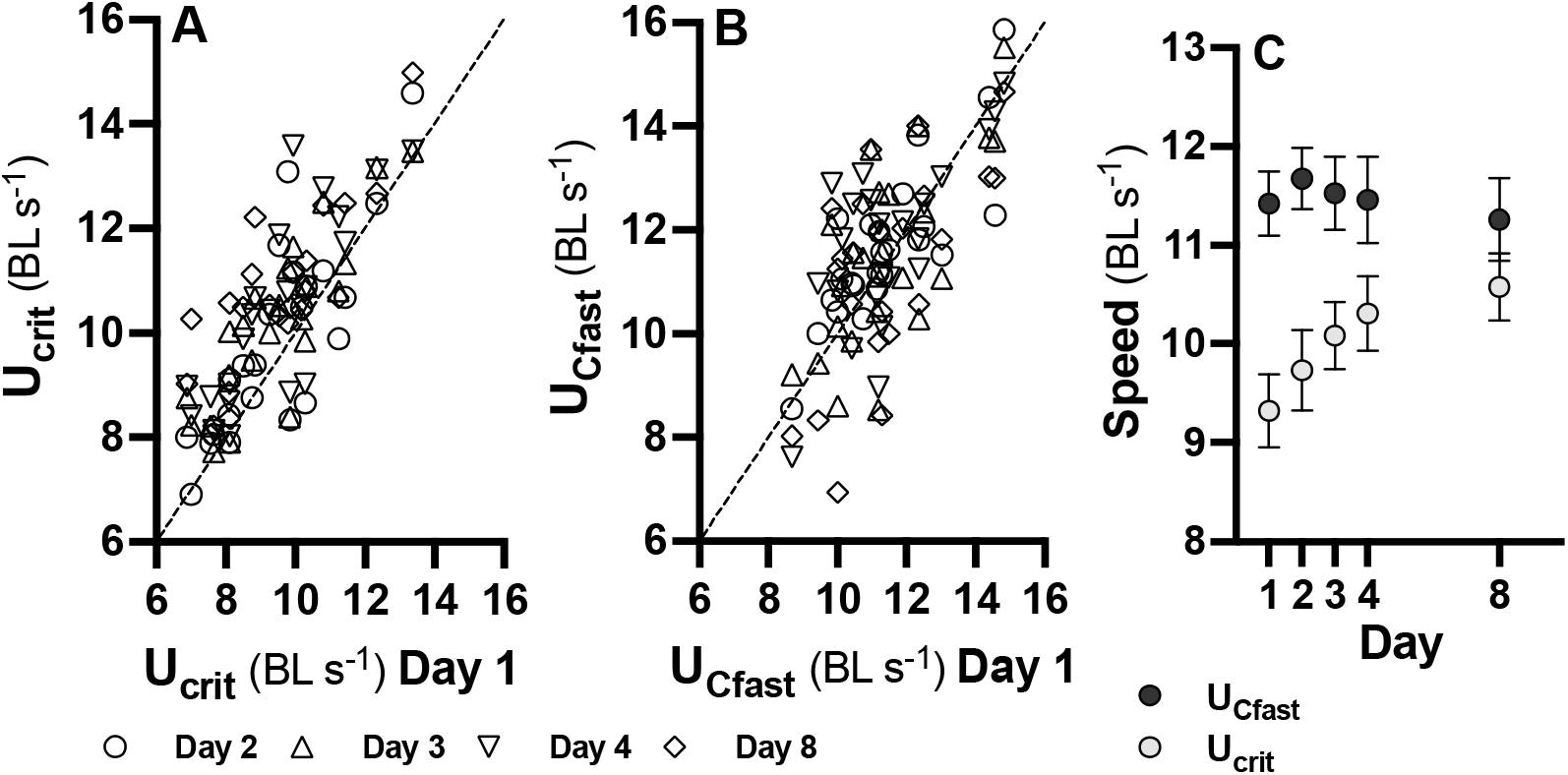
Repeatability of swimming performance. Both U_crit_ (A) and U_Cfast_ (B) were significantly repeatable (R = 0.79 and 0.69, respectively) over eight days (measurements taken on day 1 shown on x-axis, and days 2 [white circles], 3 [grey triangles], 4 [blue inverted triangles], and 8 [red diamonds] on y-axis), and repeatability did not differ between the two measures. Data from individual fish are shown. U_crit_ increased significantly with repeated swimming but U_Cfast_ did not (Fig. 1C); means ± s.e. from all fish per day are shown.

### Thermal acclimation

There were significant main effects of acclimation temperature (p < 0.001), test temperature (p < 0.001), and their interaction (p < 0.0001). The interaction indicates that 28°C acclimated fish performed better at the higher test temperature (28°C) and, similarly, that 20°C acclimated fish performed better at 20°C test temperature compared to warm acclimated fish (Fig. 2 A, B). There was a significant main effect of type of swimming performance, indicating that U_Cfast_ was overall higher that U_crit_, although the two measurements changed proportionally to each other across individuals and test temperatures (Fig. 2C, regression: R^2^ = 0.59, p < 0.001). There were no other significant interactions (all p > 0.80). Lack of significant interactions between types of swimming performance and any other fixed factors indicates that the two measures provide similar insights into thermal acclimation. U_crit_ or U_Cfast_ also gave similar individual acclimation capacity estimated derived from the double acclimation experiment (p = 0.12; Fig. 2D). The three very low values in the U_crit_ data set reduced the mean acclimation capacity, which otherwise would have been identical to that from the U_Cfast_ data.

**Figure 2.**
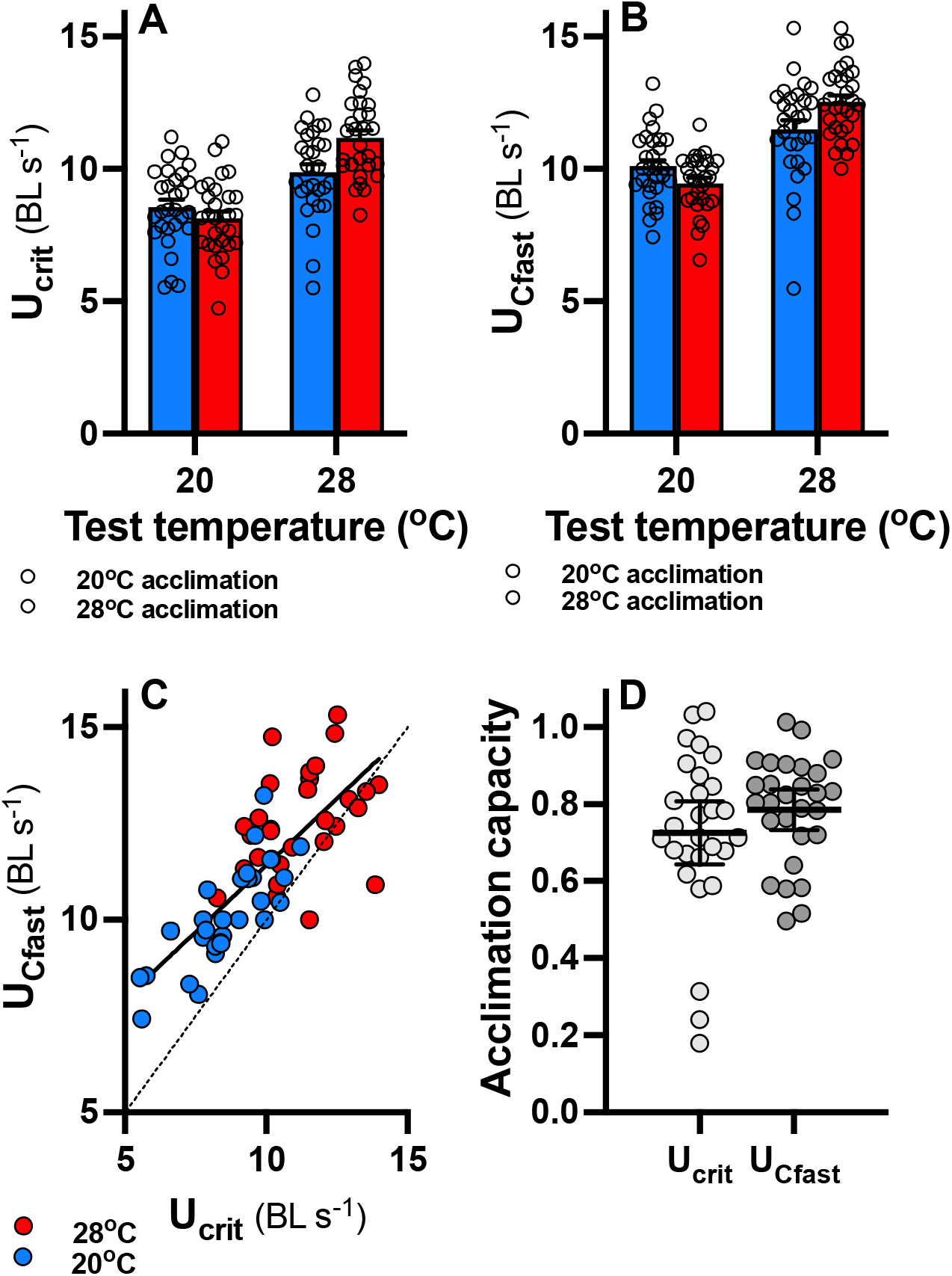
Thermal acclimation of swimming performance. Measurements of U_crit_ (A) and U_Cfast_ (B) responded similarly to acclimation to 20°C (blue bars) and 28°C and acute test temperatures of 20 and 28°C. In both cases, there were significant interaction between acclimation and test temperatures. Bars show means ± s.e., and datapoints from individual fish are shown. U_Cfast_ changed proportionally to U_crit_ across test temperatures (C; solid line = significant regression line; broken line = line of equality), but U_Cfast_ was somewhat higher particularly at cool temperatures. Fish were acclimated twice to determine individual acclimation capacity (D), which was not significantly different when based on U_crit_ or U_Cfast_; means (wide horizontal bars) ± 95% confidence intervals as well as datapoints from individual fish are shown.

In conclusion, U_Cfast_ provides similar insights into thermal acclimation as U_crit_, and both measures are equally repeatable. U_Cfast_ may even be somewhat superior because it did not induce a training effect when implemented repeatedly. U_Cfast_ is preferable to U_crit_ particularly in cases where performance is measured repeatedly such as in time-course experiments. The shorter duration of U_Cfast_ also means that much higher throughput can be achieved in experiments, which opens the opportunity for larger sample sizes particularly in more complex factorial designs.

## Author contributions

Conceptualization: FS and SMB; Experimentation: SMB; Funding: FS; Writing, first draft: FS; Writing, editing: FS, SMB.

## Data availability

All data will be deposited in Dryad upon acceptance.

## References

Burgomaster, K. A., Howarth, K. R., Phillips, S. M., Rakobowchuk, M., MacDonald, M. J., McGee, S. L. and Gibala, M. J. (2008). Similar metabolic adaptations during exercise after low volume sprint interval and traditional endurance training in humans. J. Physiol. 586, 151–160.

Denton, R. D., Higham, T., Greenwald, K. R. and Gibbs, H. L. (2017). Locomotor endurance predicts differences in realized dispersal between sympatric sexual and unisexual salamanders. Funct. Ecol. 31, 915–926.

Domenici, P. and Blake, R. (1997). The kinematics and performance of fish fast-start swimming. J. Exp. Biol. 200, 1165–1178.

Domenici, P., Allan, B. J. M., Lefrançois, C. and McCormick, M. I. (2019). The effect of climate change on the escape kinematics and performance of fishes: implications for future predator–prey interactions. Conserv. Physiol. 7, 36–22.

Drummond, G. B. and Vowler, S. L. (2012). Different tests for a difference: how do we do research? J. Physiol. 590, 235–238.

Hammer, C. (1995). Fatigue and exercise tests with fish. Comp. Biochem. Physiol. A 112, 1–20.

Husak, J. F. and Lailvaux, S. P. (2022). Conserved and convergent mechanisms underlying performance–life-history trade-offs. J. Exp. Biol. 225, jeb243351.

Husak, J. F., Fox, S. F., Lovern, M. B. and Bussche, R. A. V. D. (2006). Faster lizards sire more offspring: sexual selection on whole-animal performance. Evolution 60, 2122– 2130.

Killen, S. S., Marras, S., Nadler, L. and Domenici, P. (2017). The role of physiological traits in assortment among and within fish shoals. Phil. Trans. R. Soc. B 372, 20160233.

Killen, S. S., Cortese, D., Cotgrove, L., Jolles, J. W., Munson, A. and Ioannou, C. C. (2021). The potential for physiological performance curves to shape environmental effects on social behavior. Front. Physiol. 12, 754719.

Kolok, A. S. (1999). Interindividual variation in the prolonged locomotor performance of ectothermic vertebrates: a comparison of fish and herpetofaunal methodologies and a brief review of the recent fish literature. Can. J. Fish. Aquat. Sci. 56, 700–710.

Le Galliard, J.-F., Clobert, J. and Ferrière, R. (2004). Physical performance and darwinian fitness in lizards. Nature 432, 502–505.

López-Olmeda, J. F. and Sánchez-Vázquez, F. J. (2011). Thermal biology of zebrafish (Danio rerio). J. Therm. Biol. 36, 91–104.

Ludbrook, J. and Dudley, H. (1998). Why permutation tests are superior to t and F tests in biomedical research. Amer. Stat. 52, 127–132.

Marras, S., Peck, M., Killen, S. S., Domenici, P., Claireaux, G. and McKenzie, D. J. (2013). Relationships among traits of aerobic and anaerobic swimming performance in individual european sea bass Dicentrarchus labrax. PLoS ONE 8, e72815–12.

Plaut, I. (2001). Critical swimming speed: its ecological relevance. Comp. Biochem. Physiol. A 131, 41–50.

Seebacher, F. and Bamford, S. M. (2024). Warming and pollution interact to alter energy transfer efficiency, performance and fitness across generations in zebrafish (Danio rerio). Sci. Total Environ. 168942.

Seebacher, F., Ducret, V., Little, A. G. and Adriaenssens, B. (2015). Generalist– specialist trade-off during thermal acclimation. R. Soc. Op. Sci. 2, 140251.

Simmonds, A. I. M. and Seebacher, F. (2017). Histone deacetylase activity modulates exercise-induced skeletal muscle plasticity in zebrafish (Danio rerio). Am. J. Physiol. Regul. Integr. Comp. Physiol. 313, R35–R43.

Stoffel, M. A., Goslee, S., Nakagawa, S. and Schielzeth, H. (2017). rptR: repeatability estimation and variance decomposition by generalized linear mixed-effects models. Methods Ecol. Evol. 67, 1–6.

Svendsen, J. C., Tudorache, C., Jordan, A. D., Steffensen, J. F., Aarestrup, K. and Domenici, P. (2010). Partition of aerobic and anaerobic swimming costs related to gait transitions in a labriform swimmer. J. Exp. Biol. 213, 2177–2183.

Thambithurai, D., Crespel, A., Norin, T., Rácz, A., Lindström, J., Parsons, K. J. and Killen, S. S. (2019). Hypoxia alters vulnerability to capture and the potential for trait-based selection in a scaled-down trawl fishery. Conserv. Physiol. 7, coz082.

Wheeler, R. E. and Torchiano, M. (2016). Permutation tests for linear models in R. lmPerm version 2.1.0.

Wu, N. C. and Seebacher, F. (2022). Physiology can predict animal activity, exploration, and dispersal. Commun. Biol. 5, 109.

Zhang, Y., Claireaux, G., Takle, H., Jørgensen, S. M. and Farrell, A. P. (2018). A three-phase excess post-exercise oxygen consumption in Atlantic salmon Salmo salar and its response to exercise training. J Fish Biol 92, 1385–1403.

